# A Freeloader?: The Highly Eroded Yet large Genome of the *Serratia symbiotica* symbiont of *Cinara strobi*

**DOI:** 10.1101/305458

**Authors:** Alejandro Manzano-Marín, Armelle Coeur d’acier, Anne-Laure Clamens, Céline Orvain, Corinne Cruaud, Valérie Barbe, Emmanuelle Jousselin

## Abstract

Genome reduction is pervasive among maternally-inherited bacterial endosymbionts. This genome reduction can eventually lead to serious deterioration of essential metabolic pathways, thus rendering an obligate endosymbiont unable to provide essential nutrients to its host. This loss of essential pathways can lead to either symbiont complementation (sharing of the nutrient production with a novel co-obligate symbiont) or symbiont replacement (complete takeover of nutrient production by the novel symbiont). However, the process by which these two evolutionary events happen remains somewhat enigmatic by the lack of examples of intermediate stages of this process. *Cinara* aphids (Hemiptera: Aphididae) typically harbour two obligate bacterial symbionts: *Buchnera* and *Serratia symbiotica.* However, the latter has been replaced by different bacterial taxa in specific lineages, and thus species within this aphid lineage could provide important clues into the process of symbiont replacement. In the present study, using 16S rRNA high-throughput amplicon sequencing, we determined that the aphid *Cinara strobi* harbours not two, but three fixed bacterial symbionts: *Buchnera aphidicola,* a *Sodalis* sp., and *S. symbiotica.* Through genome assembly and genome-based metabolic inference, we have found that only the first two symbionts (*Buchnera* and *Sodalis*) actually contribute to the hosts’ supply of essential nutrients while *S. symbiotica* has become unable to contribute towards this task. We found that *S. symbiotica* has a rather large and highly eroded genome which codes only for a few proteins and displays extensive pseudogenisation. Thus, we propose an ongoing symbiont replacement within *C. strobi,* in which a once ‘‘competent” *S. symbiotica* does no longer contribute towards the beneficial association. These results suggest that in dual symbiotic systems, when a substitute co-symbiont is available, genome deterioration can precede genome reduction and a symbiont can be maintained despite the apparent lack of benefit to its host.

## Introduction

Many insects with a nutrient restricted diet, depend on vertically-inherited obligate nutritional symbionts^1–8^. These symbionts evolved from once free-living bacterial lineages^9–11^ and have undergone a series of genomic and phenotypic changes resulting from relaxed selection, continuous bottlenecks, and their metabolic adaptation to the sustained association with their host^12–15^. These alterations include genome reduction, a simplified metabolism specialised on supplying the host with essential nutrients lacking from its diet, drastic changes in cellular shape, and G+C (uncommon) or A+T-biased genomes.

Aphids (Hemiptera: Aphididae) generally house the obligate vertically-transmitted endosymbiotic bacterium *Buchnera* in specialised cells called bacteriocytes^16–18^. This obligate symbiont is capable of producing essential amino acids (hereafter **EAAs**) and B vitamins^19–24^ that are lacking from the host diet (plant phloem)^20,25,26^, and thus insures the correct development of its host^20,27,29^. *Buchnera* underwent a massive genome reduction and established as an obligate symbiont before the diversification of aphids. This is evidenced by its almost-universal presence in aphids^16,30^, the high degree of genome synteny displayed among distantly-related strains of Buchnera^31,32^, and their consistently small genomes. Aphid species from the Lachninae subfamily have been found to harbour *Buchnera* strains that have ancestrally lost the capacity to synthesise biotin and riboflavin^33–36^, two essential B vitamins. For the provision of these nutrients, Lachninae aphids and their *Buchnera* now rely on different co-obligate endosymbionts, most often *S. symbiotica*^35–38^. Accordingly, *Cinara* species (Aphididae: Lachninae) have been consistently found to host an additional bacterial co-obligate symbiont, most commonly *S. symbiotica*^38–41^. An ancestral reconstruction of the symbiotic associations of *Cinara* with fixed additional symbionts suggests that *S. symbiotica* was likely the original co-obligate endosymbiont, but has been replaced several times by other bacterial taxa^41^. These new symbionts are phylogenetically affiliated to different lineages, mainly including known aphid facultative symbiotic ones (e.g. *Fukatsuia, Sodalis,* and *Hamiltonella).*

Most of our current knowledge from these co-obligate endosymbionts comes from *S. symbiotica* strains harboured by Lachninae aphids. These symbionts display very different genomic features, ranging from strains holding rather large genomes rich in mobile elements to small genomes rich in A+T and deprived of mobile elements^42^. The *S. symbiotica* strain held by the aphid *Cinara tujafilina* (hereafter **SsCt**), shares a considerable genomic similarity to a facultative strain harboured by the pea aphid *Acyrthosiphon pisum* (hereafter **SsAp**)^35^. This reflects the early stage of genome reduction SsCt is at, which is characterised by a moderately reduced and highly rearranged genome (when compared to free-living relatives), an enrichment of mobile elements (hereafter **MEs**), and a large-scale pseudogenisation^35,43–47^. On the other side, the co-obligate *S. symbiotica* from *Tuberolachnus salignus* (hereafter **SsTs**) shows a very small and gene dense genome^36^, similarly to ancient obligate endosymbionts such as Buchnera^31–33,48,49^, *Blochmannia*^50–52^, or *Blattabacteriu*m^53–58^. Sitting in between SsCt and SsTs, the *S. symbiotica* strain housed by *Cinara cedri* (hereafter SsCc) shows intermediate characteristics between a larger and highly-pseudogenised genome and a small and compact one^37^. In *Cinara* aphids, *S. symbiotica* has undergone symbiont replacement in different lineages, and thus the endosymbionts’ genomes of species within this genus could provide important clues into reductive genome evolution and the process of symbiont replacement.

In a global survey of endosymbionts associated with about 100 Cinara species, using 16 rRNA gene high-throughput sequencing, Meseguer et al. found that Sodalis and Serratia were fixed across different populations of C. strobi.

Within the aphid *Cinara strobi,* Jousselin *et. al.* 59 first reported the presence of *Sodalis, Wolbachia,* and *Serratia* bacteria as putative secondary symbionts present in one population of this aphid species using 16S rRNA high-throughput amplicon sequencing. Later, a deeper survey of endosymbionts associated with about 100 Cinara species, using this same technique, showed that only two of these additional symbionts, *Sodalis* and *S. symbiotica,* were actually fixed across different populations of *C. strobi*^41^. *Sodalis* was found to be very abundant in both the amplicon sequencing read set and the whole-genome one. On the other hand, *S. symbiotica* was found consistently in a lower percentage than *Sodalis* in all but one sample, and was even found to be almost absent (i.e. represented by very few reads) in one (thus leading to its characterisation as a non-fixed symbiont). Further analysis of the riboflavin- and biotin-related biosynthetic genes revealed that *Sodalis* was able to supplement the previously identified auxotrophies developed by *Buchnera* strains from Lachninae aphids. This suggested that *C. strobi* most likely represented a case of co-obligate symbiont replacement, in which the former *S. symbiotica* was replaced by a younger *Sodalis* symbiont. However, this results left one unanswered question: What role, if any, is played by the prevalent *S. symbiotica* strain? We hypothesised that this bacterium could either represent a widely-spread facultative lineage (probably resembling SsAp), a transitional state in the symbiont replacement process, or a persistent *S. symbiotica* strain associated with the ancestor of *C. strobi* that had established a tripartite mutualistic symbiotic association.

To explore this question, we characterised the symbiotic community of additional populations of *C. strobi* and defined the fixed bacterial associates of this species. In addition, we assembled the genome of *S. symbiotica* from this aphid species and evaluated the metabolic capacity of its fixed symbiotic cohort to supply the aphid with EAAs, B vitamins, and other cofactors. Our results suggest that *C. strobi* houses an ancient, now dispensable, *S. symbiotica* secondary symbiont along with a co-obligate symbiotic consortium made up of *Buchnera* and its new partner, *Sodalis.*

## Materials and Methods

### Aphid collection, DNA extraction, and sequencing

*C. strobi* individuals were collected in 2015 from five colonies throughout the South Eastern Canada (supplementary table S1 in supplementary file S1, Supplementary Material online) and then kept in 70% ethanol at 6°C.

For 16S amplicon sequencing, individual aphids from each collected population (3618, 3628, 3629, 3632, and 3682) were washed three times in ultrapure water and total genomic DNA was extracted with the DNEasy Blood & Tissue Kit (Qiagen, Germany), according to the manufacturer’s recommendations. The recovered DNA was then eluted in 70 μL of ultrapure water. We amplified a 251 bp portion of the V4 region of the 16SrRNA gene^60^, using universal primers, and performed targeted sequencing of indexed bacterial fragments on a MiSeq (Illumina) platform^61^, following the protocol described in Jousselin *et. al*.^59^.

For whole-genome sequencing, we prepared DNA samples enriched with bacteria from previously collected colony 3249 following a slightly modified version of the protocol by Charles and Ishikawa62 as described in Jousselin *et. al.*^59^. For this filtration protocol 15 aphids for one colony were pooled together. Extracted DNA was used to prepare 2 custom paired-end libraries in France Génomique. Briefly, 5ng of genomic DNA were sonicated using the E220 Covaris instrument (Covaris, USA). Fragments were end-repaired, 3’-adenylated, and NEXTflex PCR free barcodes adapters (Bioo Scientific, USA) were added by using NEBNext^®^ Ultra II DNA library prep kit for Illumina (New England Biolabs, USA). Ligation products were were purified by Ampure XP (Beckman Coulter, USA) and DNA fragments (>200 bp) were PCR-amplified (2 PCR reactions, 12 cycles) using Illumina adapter-specific primers and NEBNext^®^ Ultra IIQ5 Master Mix (NEB). After library profile analysis by Agilent 2100 Bioanalyser (Agilent Technologies, USA) and qPCR quantification using the KAPA Library Quantification Kit for Illumina Libraries (Kapa Biosystems, USA), the libraries were sequenced using 251 bp paired-end reads chemistry on a HiSeq2500 Illumina sequencer. Additionally, we used reads recovered from paired-end Illumina sequencing of the same colony previously reported in Meseguer *et. al.*^41^.

### 16S rRNA amplicon taxonomic assignment

We used Mothur v1.3.363 to assemble paired-end reads and filter out sequencing errors and chimeras. In brief, the overlapped paired-end reads were assembled with the make.contigs function, and the contigs exceeding 280 bp in length were excluded from further analyses. Remaining unique contigs were then aligned with the V4 portion of reference sequences from the **SILVA** database v119^64^. Sequences that did not align with the V4 fragment were excluded from further analyses. The number of reads resulting from sequencing errors was then reduced by merging rare unique sequences with frequent unique sequences with a mismatch of no more than 2 bp relative to the rare sequences (pre.cluster command in **Mothur**). We then used the **UCHIME** program65 implemented in **Mothur** to detect chimeric sequences and excluded them from the data set. Following Jousselin *et. al.*^59^, for each sequence, the number of reads per sample was transformed into percentages using an R script and used to compile a frequency table (supplementary table S2 in supplementary file S1, Supplementary Material online). We then removed individual sequences representing less than 1/1,000 of the reads in each sample. Sequences represented by such a small proportion of the reads were generally not arthropod endosymbionts and, in most cases, were not found across PCR replicates of the same sample, suggesting that they could represent contaminants or spurious sequences.

Taxonomic assignation of the remaining sequences was conducted using the **RDP classifier**^66^ with the **SILVA** database v119 and **BLASTN**^67^ (only the best hits were reported and when hits with similar scores were found a “multi-affiliation” was reported). Using these assignations and the table of sequence frequencies per sample, we plotted the bacterial composition of each sample. To simplify representation of the results, when different unique sequences were assigned to the same bacterial species (or genus), their frequencies were added.

### Genome Assembly and Annotation

Illumina reads were right-tail clipped (using a minimum quality threshold of 20) using **FASTX-Toolkit** v0.0.14 (http://hannonlab.cshl.edu/fastx_toolkit/, last accessed December 8 2017). Reads shorted than 75 after the aforementioned clipping were dropped. Additionally, **PRINSEQ v0.20.4**^68^ was used to remove reads containing undefined nucleotides as well as those left without a pair after the filtering and clipping process. The resulting reads were assembled using **SPAdes** v3.10.1^69^ with the options --only-assembler option and k-mer sizes of 33, 55, 77, 99, and 127. From the resulting contigs, those that were shorter than 200 bps were dropped. The remaining contigs were binned using results from a **BLASTX**^70^ search (best hit per contig) against a database consisting of the Pea aphid’s proteome and a selection of aphid’s symbiotic bacteria proteomes (supplementary table S3 in supplementary file S1, Supplementary Material online). When no genome was available for a certain lineage, closely related bacteria were used. The assigned contigs were manually screened using the **BLASTX** web server (searching against the nr database) to insure correct assignment. This binning process confirmed the presence of the previously reported putative co-obligate symbionts^41,59^ (*Buchnera aphidicola* and a *Sodalis* sp.) as well as other additional symbionts. One of these additional symbionts was *S. symbiotica,* for which the first-pass assembly resulted in 3 contigs assigned to this taxon, two belonging to a putative chromosome with overlapping ends and one much smaller circular contig belonging to a putative plasmid. The resulting contigs were then used as reference for read mapping and individual genome assembly using **SPAdes**, as described above, with read error correction.

The resulting genomes were annotated using a series of specialised software. First, open reading frame (**ORF**) prediction was done using prodigal, followed by functional prediction by the **BASys** web server^71^. In order to validate start codons, ribosomal binding sites were predicted using **RBSfinder**^72^. This was followed by non-coding RNA prediction using **infernal** v1.1.2^73^ (against the **Rfam** v12.3 database^74^), **tRNAscan-SE** v2.0^75^, and **ARAGORN** v1.2.36^76^. This annotation was followed by manual curation of the genes on **UGENE** v1.28.1^7^7 through on-line **BLASTX** searches of the intergenic regions as well as through **BLASTP** and **DELTA-BLAST**^78^ searches of the predicted ORFs against NCBI’s nr database. Priority for the BLAST searches was as follows: (1) against *Escherichia coli* K-12 substrain MG1655, (2) against **Yersinia pestis** CO92 or *Serratia marcescens* strain Db11 (for *S. symbiotica),* and (3) against the whole nr database. The resulting coding sequences (CDSs) were considered to be putatively functional if all essential domains for the function were found, if a literature search supported the truncated version of the protein as functional in a related organism, or if the CDS displayed truncations but retained identifiable domains (details of the literature captured in the annotation file). For *S. symbiotica*, pseudogenes were also searched based on synteny against available *S. symbiotica* strains. This prediction performed using a combination of sequence alignment (with **m-coffee**^79^) and **BLASTX** searches against the NCBI’s nr database (restricted to *Serratia* taxon ID). This allowed the identification of missed pseudogenes by the previous searches. The annotated genomes have been submitted to the European Nucleotide Archive with project number PRJEB15507 and are on queue to be accessioned. They are temporarily available in supplementary file S4 (Supplementary Material online).

### Phylogenetic Reconstruction and Rearrangement Analysis

For performing both phylogenetic inferences and analysing the genetic differences in *Serratia* from the different aphids, we first ran an orthologous protein clustering analysis using **OrthoMCL** v2.0.9^80,81^ using a set of *S. symbiotica* and closely related free-living bacterial strains (supplementary table S4 in supplementary file S1, Supplementary Material online). We then extracted the single copy-core proteins of currently available *S. symbiotica* genomes and free-living relatives for phylogenetic reconstruction (297 protein groups) and rearrangement analysis (381 protein groups). We then ran *MGR* v2.03^82^ on the latter set to infer the tree that absolutely minimizes (no heuristics) the number of rearrangements undergone among the strains.

For phylogenetic reconstruction of *S. symbiotica,* we aligned the single-copy core protein set, gene by gene, using **MAFFT** v7.220^83^ (L-INS-i algorithm). We then removed divergent and ambiguously aligned blocks using **Gblocks** v0.91b^84^ and concatenated the resulting alignments into a single one (supplementary file S2, Supplementary Material online) for following phylogenetic inference. We used the LG+I+G amino acid substitution model, which incorporates the variability of evolutionary rates across sites in the matrix estimation^85^. Bayesian phylogenetic inference was performed in **MrBayes** v3.2.5^86^ running two independent analyses with four chains each for 300,000 generations and checked for convergence. In order to alleviate long-branch attraction artefacts commonly seen in endosymbionts^11,87^, the analysis was also run in **Phylobayes** v4.1^88^ under the CAT+GTR+G (four discrete categories) (under eight independent runs) using dayhoff6-recoded concatenated amino acid alignments. Chains were run and compared using the tracecomp and bpcomp programs, and were considered converged at a maximum discrepancy of <0.3 and minimum effective size of 50. None were found to converge even after 30,000 cycles. For further exploration, additional reduced protein datasets were selected for analysis as described above: *S. marcescens+S. symbiotica* and ribosomal proteins. Phylobayes runs done with a full set of core proteins were not found to converge even after 10,000 cycles. All resulting trees were visualized and exported with **FigTree** v1.4.1 (http://tree.bio.ed.ac.uk/software/figtree/, last accessed December 8 2017) and edited in **Inkscape**.

## Results

### Fixed symbionts of *Cinara strobi*

As stated before, *Cinara strobi* is distributed throughout eastern North America^89^. We collected *C. strobi* individuals from 5 different populations from the southeast of Canada (3618, 3628, 3629, 3632, and 3682) to complete previous sampling from northeast USA (fig. 1A and supplementary table S1 in supplementary file S1, Supplementary Material online). In order to assess the presence of bacterial associates in geographically distant *C. strobi* populations, we re-analysed the four *C. strobi* samples collected in the northeast USA (3229, 3249, 3258, and 3207), and previously included in Meseguer *et. al.*^41^, as well as the newly collected individuals through 16S rRNA high-throughput sequencing (see Materials and Methods: 16S rRNA amplicon taxonomic assignment). Taxonomic assignment of the reads revealed that individuals from all populations harboured not only two symbionts, but three: *Buchnera*, *Sodalis*, and *S. symbiotica* (fig. 1*B*). It is important to note that sample 3229 showed a very low abundance of *S. symbiotica-assigned* reads, which prompted Meseguer *et. al.*^41^ to report this symbiont as not being systematically associated with *C. strobi.* In addition to the three fixed symbionts, we also confirmed the presence of other known aphid facultative symbiont taxa (i.e. *Wolbachia, Regiella,* and *Spiroplasma*) in three samples (voucher IDs 3249, 3628, and 3628).

**Figure 1.**
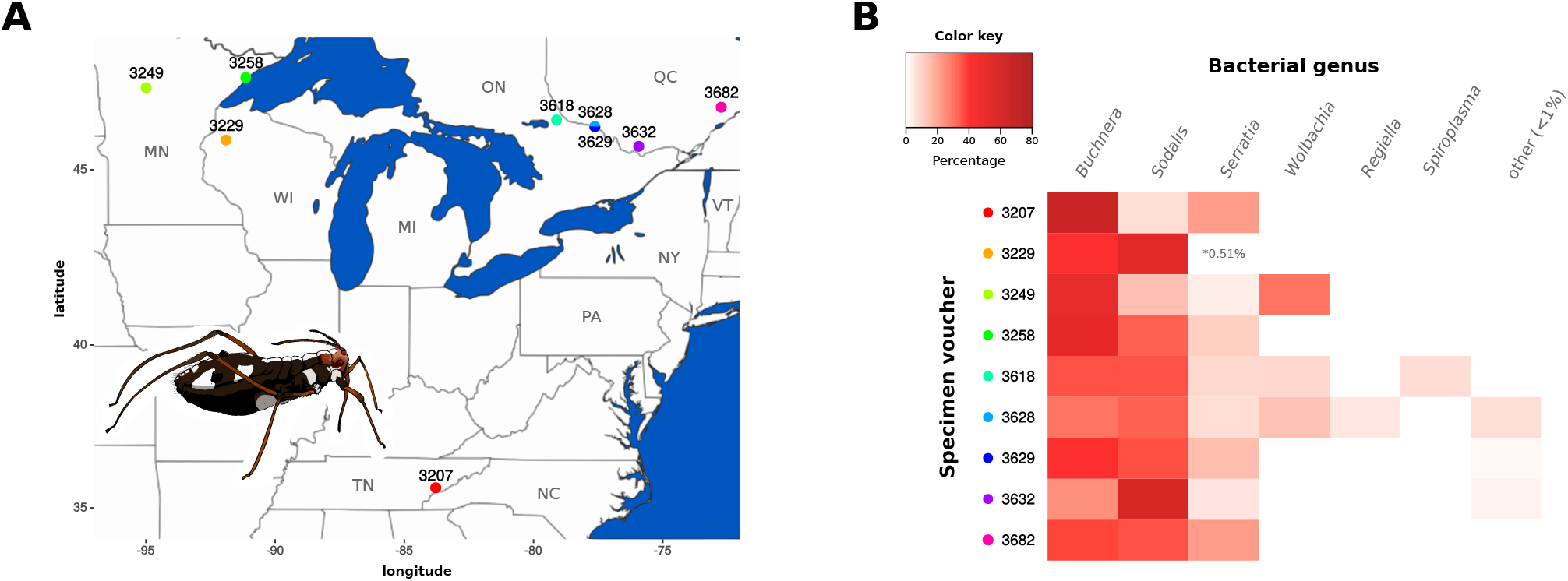
Distribution of sampled *C. strobi* populations and 16S rRNA high-throughput bacterial symbiont screening. (**A**) Map showing the north-east USA and south-east Canada regions where the *C. strobi* samples were collected (coloured points) featuring a cartoon of a *C. strobi* apterous female adult. (**B**) Heat map displaying the relative abundance of Illumina reads per taxon per sample. On the top-left, colour key for the taxon abundance. On the left, voucher ID for the sampled *C. strobi* populations with coloured dots matching the map on panel **A**.

### The genome of *S. symbiotica* strain SeCistrobi

The binning and reassembly process resulted in two assembled circular DNA molecules assigned to *S. symbiotica*: a chromosome (fig. 2A) and a plasmid, with an average coverage of 83.78x and 32.48x, respectively. The chromosome of *S. symbiotica* strain SeCistrobi (hereafter **SsCs**) is 2.41 Mbp and the plasmid is 22.67 Kbp. Its chromosome has a G+C content of 40.28%, which is slightly lower than that of both the facultative SsAp and the co-obligate SsCt (supplementary table S5 in supplementary file S1, Supplementary Material online). Unlike these two endosymbionts (which possess genomes that are similar in size), SsCs has only 635 protein coding sequences (hereafter **CDSs**), translating into a staggering low coding density of around 26.3%. This means that around 70% of its genome is non-coding, the highest known for any *S. symbiotica*. Similarly, its putative plasmid contains only two CDSs (a putative autotransporter beta-domain-containing protein and a plasmid replication protein), with the remainder of the molecule containing several pseudogenes mainly belonging to inactivated insertion sequence (hereafter **IS**) elements. Additionally, the chromosome of SsCs retains two prophage regions, however these do not encode for a single intact protein, but rather show generally highly degraded pseudogenes. Also, unlike SsAp and SsCt, it displays a typical pattern of polarised nucleotide composition in each replichore (G+C skew in fig. 2*A* and supplementary fig. S1, Supplementary Material online), hinting at a lack of recent chromosome rearrangements. This is consistent with its low number of mobile elements, when compared with SsAp and SsCt, and the complete inactivation of these by pseudogenisation and loss of other elements (e.g. inverted repeats in an IS). In regards to ncRNAs, it possesses only one rRNA operon, 38 tRNAs, a tmRNA, and 5 other non-coding RNAs (including the RNase P M1 RNA component and the 4.5S sRNA component of the Signal Recognition Particle).

**Figure 2.**
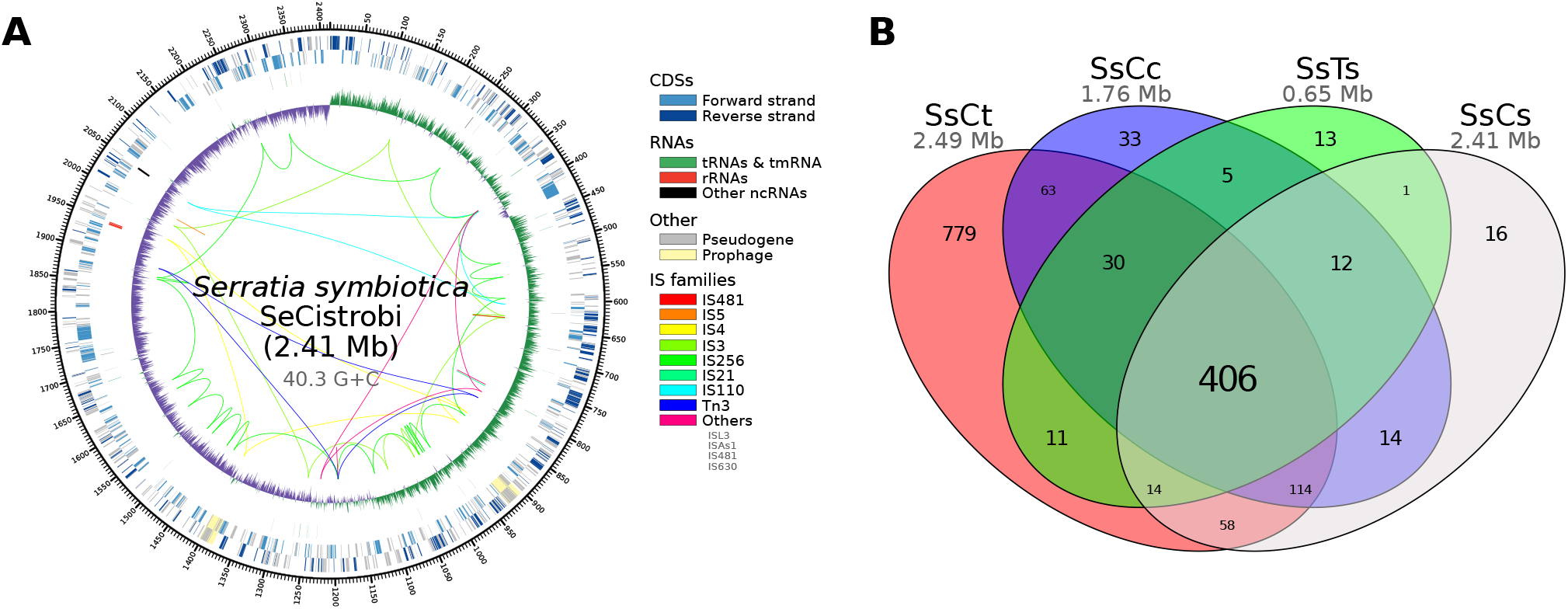
Genome of *S. symbiotica* strain SeCistrobi and pangenome of *S. symbiotica* strains from Lachninae aphids. (**A**) Genome plot of *S. symbiotica* strain SeCistrobi. From outermost to innermost, the features on the direct strand, reverse strand, ncRNA features, and G+C skew are represented. For the G+C skew, green= positive and purple= negative. (**B**) Venn-like diagram displaying the shared (core) and unshared protein-coding genes among currently-available *S. symbiotica* strains.

Regarding its CDS content, it is almost in its entirety a subset of the pan-genome of *S. marcescens*, except for the two plasmid CDSs (supplementary fig. S2, Supplementary Material online). While the putative autotransporter beta-domain-containing protein from SsCs does not cluster with any other proteins, its best 5 matches in NCBI’s nr database are against other autotransporter beta-domain-containing proteins from *S. symbiotica* strain CWBI-2.3. Therefore it shows as strain specific in our analysis due to the strains chosen for the protein clustering. When compared with co-obligate *S. symbiotica* strains from Lachninae aphids fig. 2B, it shares most of its genetic repertoire with the highly reduced SsCc and SsTs strains. Within the subset of non-core genes, we observed mainly genes retained in degraded pathways, as well as others that reflect differences in pathway retention (such as difference in the metabolism of nucleotides, gluconeogenesis, and sulfur cluster biosynthesis), and genes that code for membrane proteins both involved in the transport of different compounds (such as putrescine import, Sodium/proline symport, and arginine transport) and of unknown function. In terms or DNA repair, SsCs retains mostly the same set of proteins as the most genomically reduced *S. symbiotica* symbionts (SsCc and SsTs), with the marked exception of SsCc retaining Dam, MutH, MutL, and MutS; thus coding for a mismatch repair system lacking the exonucleases ExoX, XseA, XseB, RecJ, and the non-essential HolE protein from the DNA polymerase III.

We reconstructed phylogenetic trees using 297 single-copy CDSs that were shared by all *S. symbiotica* strains, a selection of free-living *Serratia*, and *Yersinia pestis* strain CO92 (as an outgroup). Using **MrBayes**, we found *S. symbiotica* as a monophyletic group sister to the *S. marcescens* clade (supplementary fig. S3A, Supplementary Material online). Given the very long branches leading to the highly reduced SsCs, SsCc, and SsTs; we also ran a phylogenetic reconstruction in **Phylobayes** with dayhoff-6 recoded alignments and under the CAT+GTR+G (four discrete categories) model. This method is presumably less sensitive to long branch attraction artifacts commonly seen in phylogenies including highly derived endosymbiont lineages^11,87^. From all 8 independent chains we ran, only two of them converged, even after 24,000 generations (wit some even reaching the 28,000 and 30,000 generations). However, the *S. marcescens+S. symbiotica* clade, as well as other bipartitions, were lowly supported and/or unresolved (supplementary fig. S3B, Supplementary Material online). Additional reconstructions were performed using subsets of these data with **MrBayes** and *Phylobayes* (see Materials and Methods: Phylogenetic Reconstruction and Rearrangement Analysis), finding similar results in the former and alternative topologies for the latter (file S3, Supplementary Material online). This suggests that additional taxa (e.g. *Serratia* strains associated with other *Cinara* species) are probably needed to stabilise the phylogenetic trees. Finally, and like all other currently available *S. symbiotica* strains, its genome shows many rearrangements (supplementary fig. S3C, Supplementary Material online) when compared to free-living *S. marcescens* and other *S. symbiotica*.

### Biosynthesis of Essential Amino Acids and B Vitamins by the symbiotic consortium in *Cinara strobi*

In previously analysed co-obligate endosymbiotic systems in Lachninae aphids (Buchnera+secondary symbiont), *Buchnera* remains as the sole provider of EAAs and the newly acquired symbionts have taken over the role of synthesising riboflavin (vitamin B2) and biotin (vitamin B7), functions once performed by Buchnera^35–37,41^. Thus, to infer the role of each fixed symbiont of *C. strobi,* we searched for the genes involved in the biosynthesis of EAAs (fig. 3), B vitamins, and other cofactors (fig. 4 and supplementary fig. S4, Supplementary Material online) in *Buchnera, S. symbiotica,* and *Sodalis* from *C. strobi* and compared them with co-obligate *Buchnera+Serratia* endosymbiotic systems in Lachninae (using *Buchnera-only* Aphididae systems as reference).

**Figure 3.**
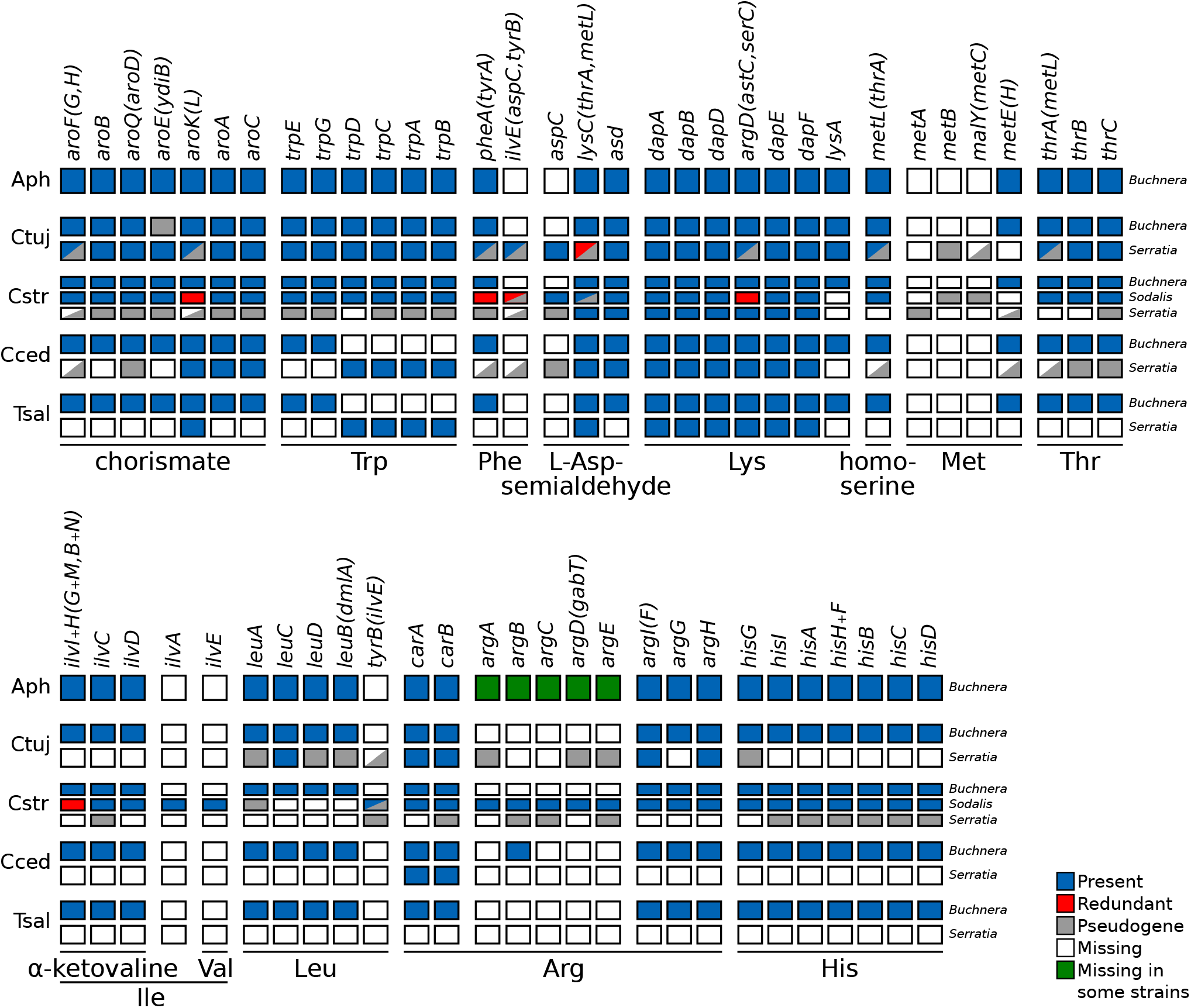
Essential-amino-acid biosynthetic metabolic capabilities of obligate symbiotic consortia of different aphid species. Diagram summarising the metabolic capabilities of the fixed endosymbiotic consortia of co-obligate symbiotic systems of Lachninae aphids. For comparison, a collapsed representation of Aphididae *Buchnera*-only systems is used as an outgroup. The names of genes coding for enzymes involved in the biosynthetic pathway are used as column names. Each row’s boxes represent the genes coded by a symbiont’s genome. At the right of each row, the genus for the corresponding symbiont. Abbreviations for the aphids harbouring the symbionts is shown at the left of each group rows and goes as follows. Aph= Aphididae, Ctuj= *C. tujafilina,* Cstr= *C. strobi,* Cced= *C. cedri,* Tsal= *T. salignus.* On the bottom, lines underlining the genes involved in the pathway leading to the compound specified by the name underneath the line. For amino acids, their three letter abbreviations are used.

**Figure 4.**
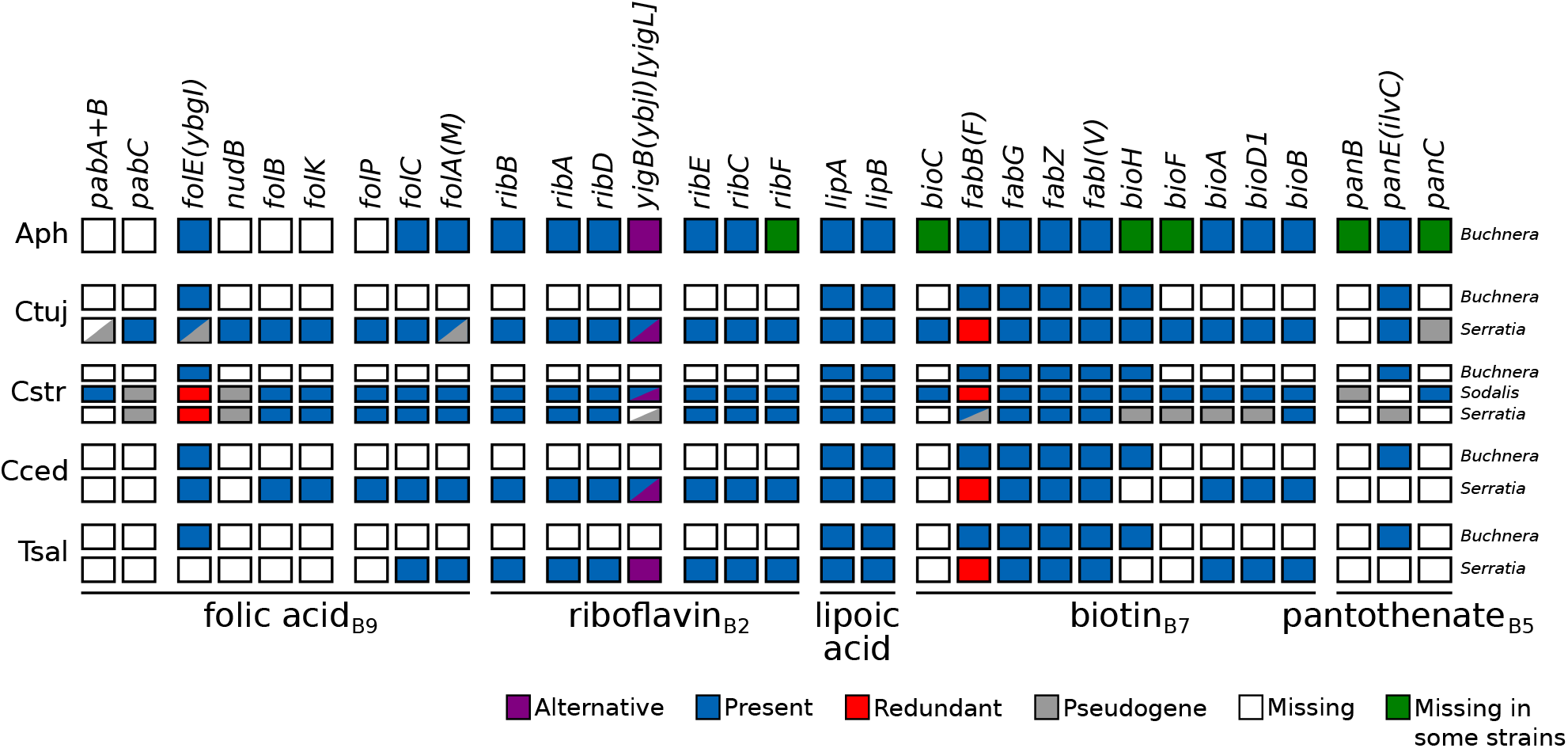
B-vitamin biosynthetic metabolic capabilities of obligate symbiotic consortia of different aphid species. Diagram summarising the metabolic capabilities of the fixed endosymbiotic consortia of co-obligate symbiotic systems of Lachninae aphids. For comparison, a collapsed representation of Aphididae *Buchnera-only* systems is used as an outgroup. The names of genes coding for enzymes involved in the biosynthetic pathway are used as column names. Each row’s boxes represent the genes coded by a symbiont’s genome. At the right of each row, the genus for the corresponding symbiont. Abbreviations for the aphids harbouring the symbionts is shown at the left of each group rows and goes as follows. Aph= Aphididae, Ctuj= *C. tujafilina,* Cstr= *C. strobi,* Cced= *C. cedri,* Tsal= *T. salignus.* On the bottom, lines underlining the genes involved in the pathway leading to the compound specified by the name underneath the line.

In terms of EAAs, *Buchnera* from *C. strobi* (hereafter **BCs**), retains the same capabilities as other *Buchnera* strains. Similarly, *Sodalis* also retains all genes needed for the biosynthesis of EAAs, except for those of lysine, methionine, and leucine. In the case of SsCs, it has completely lost the potential of *de novo* synthesising all EAAs. Nonetheless, it preserves an almost intact route for the synthesis of lysine, resembling the degradation pattern observed for this pathway in other co-obligate *S. symbiotica* strains.

Regarding B vitamins and other cofactors, we found that BCs is unable to synthesise vitamin B2 and B7, similarly to the other *Buchnera* from Lachninae aphids. Unlike the Lachninae co-obligate endosymbiotic systems, we determined that SsCs is unable to takeover the role of synthesising these two vitamins. The vitamin B2 pathway would be interrupted due to the loss of a 5-amino-6-(5-phospho-D-ribitylamino)uracil phosphatase enzyme, preserving only a *yigB* pseudogene (interrupted by various stop codons and frameshifts). From the genes needed to complement *Buchnera’s* truncated biotin pathway (*bioA, bioD1,* and *bioB*), it preserves only *bioB*. However, it still retains identifiable pseudogenes for *bioA* and *bioD1*. All other pathways for B vitamins and other cofactors are degraded, except for that of lipoic acid. On the other hand, and as previously reported by Meseguer *et. al.*^41^, *Sodalis* is indeed able to takeover the role as the provider of both riboflavin and biotin, thus being essential for the beneficial symbiosis.

## Discussion

Genome degeneration is a common characteristic of vertically-inherited mutualistic symbionts of insects^90,91^, and is particularly marked in ancient nutritional mutualistic endosymbionts^33,48,52,92^. These genome deterioration can eventually affect pathways involved in the symbiont’s essential functions, such as those involved in essential-amino-acid or B-vitamin biosynthesis. When this occurs, the symbiont is either replaced by a more capable symbiont, or is complemented by a new co-obligate symbiont^15^. As members of the Lachninae subfamily, *Cinara* aphids depend on both *Buchnera* and an additional symbiont for the supply of essential nutrients, namely EAAs and B-vitamins^35,37,41^. While *S. symbiotica* is the most prevalent and putatively ancestral symbiont, it has been replaced by other bacterial taxa in several lineages^41^. *Cinara strobi* represents such a case, in which the putatively ancient co-obligate *S. symbiotica* symbiont has been replaced by a *Sodalis* strain.

Here, we further explored the composition and the role of the fixed symbiotic cohort of the aphid *C. strobi.* Through the re-analysis of previously reported 16S rRNA NGS amplicon data from geographically distant *C. strobi* populations plus additional ones, we found that not only *Buchnera* and *Sodalis* were fixed, but also *S. symbiotica*. This third symbiont was previously not deemed as fixed given the low abundance (< 1%) of NGS amplicon reads assigned to this taxon, consistent with the low amount of whole-genome sequence data belonging to *S. symbiotica*^41^. Thus, the persistent association of this symbiont across populations of *C. strobi* points towards this being a non-facultative, hence obligate, symbiotic relationship.

Through whole-genome sequencing of the genome of SsCs, we have provided evidence that SsCs could well be a missing link between the loss of function of a symbiont and the acquisition of a new and more capable one. In spite of SsCc showing a large genome (2.41 Mbps), it displays drastic genome pseudogenisation (around 26.3% coding density). This drastically contrast both the “early” co-obligate SsCt (~2.49 Mbps and 53.4% coding density) and the “modestly” shrunk co-obligate SsCc (1.76 Mbps and 39.0% coding density^)42^. This means that the majority of SsCs’ genome is made up of pseudogenes and “genomic wastelands”. This would place the genome in an intermediate state of reduction, before losing bigger chunks of it and thus, evolving a smaller-sized genome. The evolutionary relation SsCs keeps with other *S. symbiotica* symbionts remains uncertain. Consistent with a previous phylogenetic reconstruction^36^, we found that, through the use of inference methods that alleviate long branch attraction artefacts, the relationships among *S. symbiotica* lineages is not well resolved. This could be due to the extremely long branches, seen in SsCs, SsCc, and SsTs; when compared to SsAp, SsCt, and other free-living *Serratia;* which confounds phylogenetic signal (see^87^). This makes it difficult to interpret the evolutionary origin and relationships of *S. symbiotica* endosymbionts solely from phylogenetic data. We expect further large-scale sequencing of these endosymbionts will provide further data to disentangle *S. symbiotica* phylogenetic relationships.

G+C skew in transitional genomes from some endosymbiotic lineages show an altered pattern, when compared to free-living relatives^9^ or long-term highly-reduced endosymbionts^36,52^. This perturbation may result from recent chromosome rearrangements likely due to recombination events between repetitive elements, namely ISs^9^. The presence of a typical pattern of polarised nucleotide composition in each replichore of SsCs (fig. 2A) points towards long-term genome stability, consistent with the lack of functional mobile elements. This G+C skew pattern is not observed neither in the facultative SsAp nor the co-obligate SsCt (supplementary fig. S1, Supplementary Material online). Therefore, the G+C skew pattern in SsCs, together with its highly degenerated genome and the fixed presence of *S. symbiotica* in different aphid populations, hints at both a long-term obligate association and a vertical transmission of the symbiont in *C. strobi*.

When a symbiont replacement occurs, it is expected that the new symbiont will replace the symbiotic functions of the former one. This is seen in different mono- and di-symbiotic systems observed in weevils^93^, aphids^41,94,95^, mealybugs^96,97^, and several Auchenorrhyncha^46,98,99^. As observed in all other currently sequenced *Buchnera* from Lachninae aphids, BCs is unable to provide two essential B vitamins: biotin (B7) and riboflavin (B2). In the case of *C. strobi,* Meseguer *et. al.* 41 found that *Sodalis* was capable of supplementing this deficiencies, thus making this fixed symbiont essential for both *Buchnera* and the aphid. Here, we have found that these two fixed symbionts indeed are together capable of producing all EAAs and B vitamins for their aphid host and each other. When looking at SsCs, the third fixed symbiont in *C. strobi*, we found that it is unable to independently synthesize any of the aphid’s essential nutrients. This suggests that this symbiont is no longer contributing to the co-obligate nutritional endosymbiotic consortium in *C. strobi* but it has persisted in the aphid regardless its metabolic dispensability. The retention of certain enzymes in pathways related to the synthesis of EAAs and B vitamins, can be explained in two ways: i) The enzymes have not had enough time to accumulate mutations which would render them pseudogenes, and ii) These enzymes participate in other cell-maintenance pathways. Evidence for the former is observed in other *S. symbiotica* and *Sodalis* genomes, which display various degraded pathways (theoretically inactive but still coding for several enzymes^35,45,47^. Relating to the latter, the retention of several genes in the biotin pathway *fabB, fabG, fabZ, fabV,* and *bioH*) is possibly due to their involvement in the pathway leading to cell-membrane biogenesis.

It could be argued that SsCs could be a widespread pathogen. However, the likely ancestral presence of this symbiont as a co-obligate symbiont^41^, its universal presence in the sampled populations from the species, the highly reduced genome, the lack of proteins with identifiable eukaryotic-like domains (e.g. ankyrin- or leucine-rich repeats) or diverse secretion systems, does not support this hypothesis.

The genome of SsCs also reveals that a massive genome reduction does not necessarily preclude the symbiont’s replacement. The low amount of intact CDSs that SsCs preserves could be explained by the fixation of *Sodalis* as a co-obligate symbiont. The long-term association with this new symbiont would thus relax selective pressure on keeping a number of genes, namely those that are redundant. This pattern of gene loss following the acquisition of a companion symbiont can be seen in at least two co-obligate systems: Buchnera+secondary in aphids^36^, and Tremblaya+secondary in mealybugs^96^ (see^15^). It is worth noting the retention of a mismatch repair system in SsCs, which is involved in the detection of non-Watson-Crick base pairs and strand misalignments arising during DNA replication^100^. However, the retention of this system does not, to our knowledge, help explain the retention of a large genome with such a low coding capacity. This retention could rather partly explain the lack of an extreme A+T-biased genome (see14), such as the ones held by SsCc and SsTs.

Taken together, the evidence points towards a di-symbiotic co-obligate system in *C. strobi,* with the two co-obligate partners being *Buchnera* and *Sodalis.* Based on an ancestral reconstruction of symbiotic associations in in *Cinara*^41^, this case would constitute one of secondary co-obligate symbiont replacement. At some point in the lineage of *C. strobi,* the putative ancient secondary co-obligate *S. symbiotica* symbiont would have been metabolically replaced by the new and capable *Sodalis.* Whether the inactivation of the genes involved in the *de novo* synthesis of both riboflavin and biotin happened before the acquisition of *Sodalis* (rescue) of after it (takeover) through relaxed selection on the retention of those genes, remains unclear. Following this loss of symbiotic function, *S. symbiotica* would have continued to thrive within the aphid and be vertically inherited from mother to offspring. The perpetuation of *S. symbiotica* in *C. strobi* could hypothetically be a collateral result of a fine-tuned system of symbiont inheritance in the aphid. A similar case could be made for *Westeberhardia,* the putative ancient endosymbiont of at least some *Cardiocondyla* ants^101^. In *Cardiocondyla obscurior,* the symbiont inhabits the cytoplasm of bacteriocytes and possesses a very small genome (532.68 kbp). Its genome lacks intact pathways for the biosynthesis of any EAA or B vitamin, but codes for 4-hydroxyphenylpyruvate. This last can be converted into tyrosine by the ant host, thus the symbiont would hypothetically contribute to cuticle formation during the pupal stage. Interestingly, the authors report on a natural population that has lost this symbiont and seems to thrive in the laboratory (at least under conditions including *ad libitum* protein provisioning). This reflects *Westeberhardia* has possibly been retained in other populations despite its apparent dispensability. Thus, the loss of an otherwise long-term symbiont like SsCs would require mutational loss of it and subsequent fixation through drift.

## Conclusion

Based on the genome-based metabolic analysis of the pathways involved in the synthesis of EAAs and B-vitamins, we have found that only *Buchnera* and *Sodalis* are required for the provision of these nutrients to the aphid. *S. symbiotica,* the third fixed symbiotic partner, does not seem to be contributing towards the mutualistic consortium, suggesting that it has effectively become a “freeloader” which likely evolved from an ancient co-obligate lineage. Our results reveal that after an obligate symbiont’s metabolic-based replacement, the formerly essential associate can be perpetuated in a consortium despite its dispensability. Also, the genome of SsCs evidences that a long-term symbiont can retain a rather large genome despite its extreme low coding density. We expect the exploration of other *Buchnera*+*S. symbiotica* co-obligate systems from closely related lineages to *C. strobi* will further illuminate the genome reduction process undergone by this symbiont as well as the reasons behind its overstay as a “freeloader” in this aphid species.

## Acknowledgements

We would like to acknowledge the talented artist/scientist Jorge Mariano Collantes Alegre for the aphid cartoon in fig. 1A. This work was supported by the Marie-Curie AgreenSkills+ fellowship programme co-funded by the EU’s Seventh Framework Programme (FP7-609398) to A.M.M., the Agropolis foundation/Labex Agro (“Cinara’s microbiome”) to E.J, the the *France Genomique* National Infrastructure, funded as part of the *Investissemnt d’Avenir* program managed by the *Agence Nationale pour la Recherche* (ANR-10-INBS-09) to C.O, C.C., and V.B. This publication has been written with the support of the AgreenSkills+ fellowship programme which has received funding from the EU’s Seventh Framework Programme under grant agreement No. FP7-609398 (AgreenSkills+ contract). We are grateful to the genotoul bioinformatics platform Toulouse Midi-Pyrenees (Bioinfo Genotoul) for providing help and/or computing and/or storage resources. The authors are grateful to the CBGP-HPC computational platform. The funders had no role in study design, data collection and analysis, decision to publish, or preparation of the manuscript.

